# Inbreeding or outbreeding depression? How to manage an endangered and locally adapted population of red grouse *Lagopus scotica*

**DOI:** 10.1101/2023.08.14.552414

**Authors:** Grace Walsh, Barry J. McMahon, Filip Thörn, Patrik Rödin-Mörch, Martin Irestedt, Jacob Höglund

**Author notes:** Corresponding author: Grace Walsh.

## Abstract

The balance between the inbreeding and outbreeding continuum is a fundamental debate within conservation genomics. To inform this debate we use genome wide data to show that inbreeding is higher in Irish red grouse (*Lagopus scotica*) and that this smaller and isolated population exhibits divergence from the British. Outlier analysis identified divergence in several genomic regions linked to genes coding for ecologically important traits such as phenotypic differences in plumage colour and sequence divergence among coding regions in the melanin pathway. We also find differences among immune genes and loci involved in food intake and energy control. The two populations thus appear locally adapted. Divergence between the source and the target population when used for conservation aided restocking may have negative consequences if locally adapted alleles are swamped and/or maladapted genotypes are introduced, leading to outbreeding depression. While avoiding inbreeding by maintaining large populations is important, our study illustrates that conservationists and managers should also consider population divergence and local adaptation. We argue against translocations among Ireland and Britain as a conservation strategy. The results of our study demonstrate the value of focusing on local populations and habitat management when aiming to enhance the conservation status of specific species.

## Introduction

In contemporary conservation science there is an ongoing debate on how to manage small and isolated populations (Hohenlohe et al., 2021; Kardos et al., 2021). Some researchers advocate that the most serious threat to population survival is inbreeding depression caused by an accumulation of realized genetic load. Inbreeding is exacerbated in small and isolated population fragments and the suggested management strategy is to avoid introducing masked load when translocations from large to small populations when attempts to save a smaller population are implemented (Grossen et al., 2020; Robinson et al., 2019, 2018; Teixeira and Huber, 2021). Such a strategy would recommend that managers should be careful when moving individuals among population fragments. An alternative school advocate extensive translocation in order to increase genetic variation in small and genetically depauperate populations (Allendorf, 2017; Whiteley et al., 2015). The latter strategy has been termed “genetic rescue” (Frankham, 2017). While there seems to be consensus that some translocations should be undertaken at least when population size of the fragmented population is very small and the genetic diversity is low, managers have to consider whether or not the translocations may introduce genetic load and maladapted gene complexes to the population aimed to be rescued (Kyriazis et al., 2021; van Oosterhout, 2020).

The red grouse is an iconic bird species recently elevated to species status (Sangster et al., 2022) found in the United Kingdom and Ireland. The Irish population is smaller by an order of magnitude than the British (ca. 4000 vs. 250.000 breeding pairs (Allen et al., 2005; Cummins et al., 2015; Woodward et al., 2020)) and is considered a conservation concern in Ireland (Cummins et al., 2015; National Red Grouse Steering Commitee, 2013). Previous research has shown that Irish and British red grouse are genetically differentiated at neutral (McMahon et al., 2012; Meyer-Lucht et al., 2016) and adaptive markers (Meyer-Lucht et al., 2016). Studies have also indicated a significant reduction in the genetic diversity of Irish red grouse (Freeland et al., 2007),. The question then arises, are translocations from Great Britain to Ireland a viable conservation strategy?

Here we use genome wide data to show that the smaller and isolated population in Ireland show divergence to the British in a number of genomic regions that can be linked to genes coding for ecologically important traits. We show the plumage of Irish grouse is phenotypically paler than British and that the populations are divergent in a number of genes involved in feather development and the melanin pathway determining pigmentation. We also find evidence for divergence among immune genes, genes involved in feeding efficiency and other genes implicated in coding for other ecologically important traits. The two populations thus appear locally adapted to their respective island. We argue against translocations between Ireland and Great Britain as it may cause outbreeding depression. Instead, we advocate that further fragmentation of Irish population should be avoided and local population sizes and measures augmenting within Ireland dispersal should be implemented. Our study illustrates that while avoiding inbreeding depression by maintaining large enough populations sizes, managers also need to consider the divergence between the source from where assisted dispersal and translocations are taken and the target population aimed to be rescued.

## Results

### Sequences, Mapping, SNP calling and Genetic Diversity

After mapping and filtering the reads the mean depth per sample was 24.3x. Details of these samples are shown in Table S1. The final SNP dataset with all individuals consisted of 3345069 SNPs. The dataset which was pruned for linkage disequilibrium contained 662871 SNPs. The final SNP dataset for only Recent Irish and English individuals contained 5873882 SNPs of which 849131 remained after pruning for linkage disequilibrium. Nucleotide diversity (π) was similar in all populations. It was lowest in the museum samples at 0.00088 (± 0.00059 SD) and very similar in the English and Recent Irish samples at 0.00116 (± 0.00071 SD) and 0.00107 (± 0.00067 SD), respectively.

### Principal component analysis

A PCA showed genetic structure corresponding to the sample locations. Together PC1 and PC2 explained 20.33% of the variance as shown in Fig. 1b. The Irish and English samples split along PC1 but not PC2. The analysis also shows the English samples grouping more closely together than the Irish samples, which corresponds to the geographical extent of sampling in both countries (see Fig. 1a).

**Figure 1.**
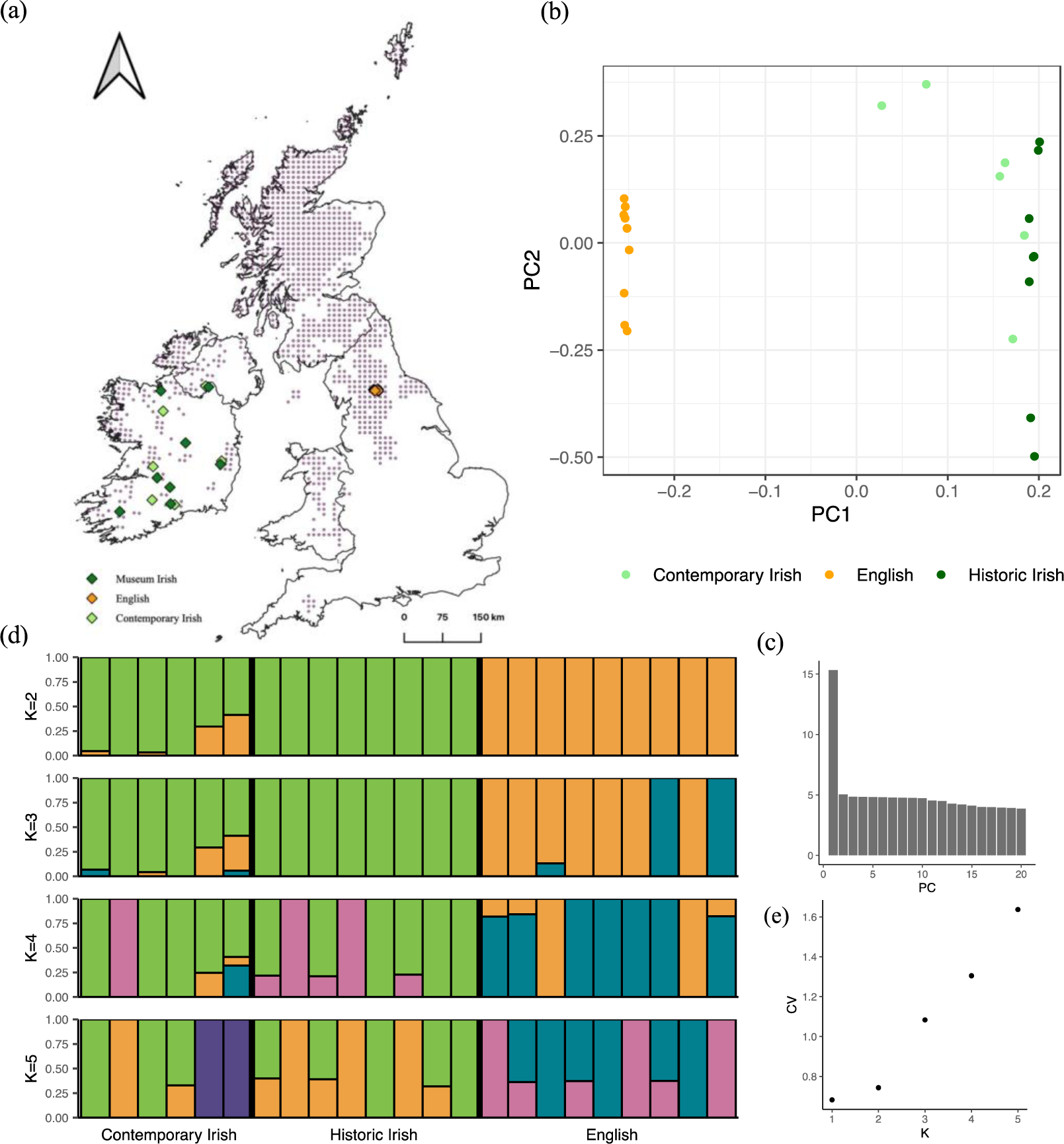
(**a**) Map of the British Isles showing the year-round distribution of red grouse. Distribution is shown on a 10km grid square level which each dot representing a grid square where red grouse were detected. English samples are shown in red, Recent Irish samples (between 2007-2020) are shown in orange and Museum samples (c.1880) are shown in purple. Map reproduced from Bird Atlas 2007–11, which is a joint project between, BTO, BirdWatch Ireland and the Scottish Ornithologists’ Club. Map reproduced with permission from the British Trust for Ornithology. (**b**) Principal component analysis (PCA) of genetic variation for 23 individuals of red grouse from Ireland and England. Each dot represents one individual with orange representing English birds and blue representing Irish birds. **(c)** Bar chart showing the amount of variation explained by the first 20 principal components. **(d)** Admixture analysis performed with ADMIXTURE for K=2-5 genetic clusters. Each vertical bar represents one individual and the colours group them by genetic cluster. The solid black lines separate the three different populations of Contemporary Irish (n=6), Historical Irish (n=8) and English (n=9). The y-axis shows the ancestral fractions. Bars of a single colour imply ancestry from a single population. **(e)** Cross-validation error for ADMIXTURE runs with K set to 1-5.

PC3 and PC4 explained c. 5% variation each as shown in Fig. S1 (b). When the effect of population (PC1) is removed the Irish samples cluster closely together and the English samples are spread out Fig. S1 (b). This suggests that when the effect of location is removed the Irish samples are less genetically diverged than the English sample.

### Microsatellites

Microsatellite allelic variation at 15 neutral microsatellite loci were scored for 7 birds from Scotland (Aberdeenshire) and 15 birds sampled in Northern Ireland. These were the same 15 loci previously typed in other populations throughout Ireland(Johansson et al., 2012; McMahon et al., 2012). Expected multi-locus heterozygosity (± SD) among the samples from Northern Ireland was 0.695 ± 0.036 while observed heterozygosity was 0.546 ± 0.036.

A DAPC-plot of the microsatellite variation of the birds from Northern Ireland with birds from the previously reported sampling sites in the Republic of Ireland (McMahon et al., 2012) revealed birds from Northern Ireland were separated from birds from the Republic along the first axis (Fig. S2). The global multilocus FST was 0.03 revealing a low but highly significant population structure (P <0.0001). Pair-wise estimates reveal that Northern Ireland was significantly different from all the other sites (except birds sampled in the west of Ireland counties Mayo and Galway, however sample size was low n=3 in in this region) (Table 1). When we also added seven birds from Aberdeenshire, Scotland to a DAPC, birds from Northern Ireland fell in-between the Scottish sample and the samples from the Republic of Ireland. This comparison revealed a highly significant population structure (FST=0.03, P <0.0001, Fig S2).

**Table 1.** Pair-wise FST estimates for 15 microsatellite loci. Significant values in bold. Note that the sample size for population “West” was only 3 birds.

|  | **Cork** | **Munster** | **N. West** | **N. Ireland** | **Wicklow** |
| --- | --- | --- | --- | --- | --- |
| **Munster** | 0.0395 |  |  |  |  |
| **N. West** | 0.0329 | 0.0207 |  |  |  |
| **N. Ireland** | 0.0410 | 0.0249 | 0.0343 |  |  |
| **Wicklow** | 0.0342 | 0.0202 | 0.0145 | 0.0465 |  |
| **West** | 0.0284 | 0.0032 | -0.0364 | -0.0030 | 0.0083 |

### Analysis of admixture

The lowest cross validation value, except for k=1 was for k=2 in the ADMIXTURE analysis. This assigned populations to genetics clustered which corresponded with the geographic locations of the samples (Fig. 1e). Fig. 1d shows the admixture results for 2-5 populations. When k is set to 2 it shows all English samples forming one distinct population. Of the 14 Irish individuals, two contemporary individuals show admixture of c. 30-40% and another two contemporary individuals show minor admixture from the English population. The admixed individuals still show predominant ancestry from the Irish population.

### Window-based FST scans

The mean FST between the Recent Irish and English samples was 0.095. The spatial distribution of ZFST across the genome (Fig. 2) shows several peaks of differentiation. The major peaks are present on chromosomes 3, 4, 6 and 7. The windows with the highest ZFST scores are on chromosome 3 at 11.3 and 11.1 followed by chromosome 6 with two windows of 10.65 ZFST. The distribution of ZFST along the genome does not appear to be uniform.

**Figure 2.**
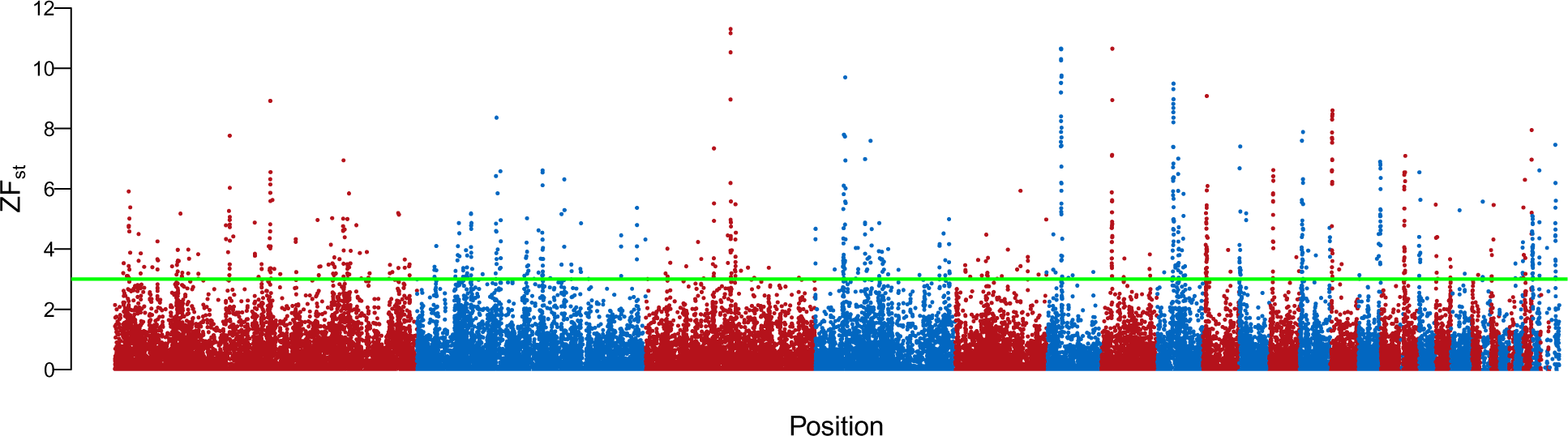
Manhattan plot of genome-wide F_ST_ outlier scan for autosomal chromosomes from Recent Irish vs. English birds. The y-axis shows ZF_ST_. A score >3 (indicated above the green line) was considered an outlier. Each colour change indicates a changing chromosome. Each dot represents the average ZF_ST_ value for a 15 kb window.

### Outlier Analysis

We used two methods used to identify outlier candidate genes associated with local adaptation. Outlier regions were mapped to the annotated chicken genome to extract genes within them. First, there was a total of 12016 outlier SNPs found using the pcadapt method on contemporary samples. These SNPs were not located within any coding genes. Within a 5kb region upstream and downstream there were 545 genes extracted and when duplicates were removed 333 remained. These SNPs are likely located in promotor regions or transcription factors and therefore could have functions relating to the nearby genes. Second, window-based FST scans were also used to detect outliers. FST values were transformed to ZFST. The genes in any windows with ZFST >3 were extracted. This method identified 345 genes within 5kb upstream and downstream of the 15kb outlier regions. This resulted in a total list of 661 outlier genes. Of these, 16 genes were identified by both methods (Table S2). The full lists of genes for ZFST outlier method are shown in Table S3 and Table S4 shows the full list of outliers using the pcadapt method.

### Phenotypic colour differences and candidate pigmentation genes

Irish birds reflected more light from the light bars of the body feathers than either British or Scandinavian grouse (Kruskal-Wallis χ^2^ =19.77, df=2, P < 0.0001, Fig. 3a). There were no significant differences among British and Scandinavian birds (P > 0.10) and no differences among the regions with respect to reflectance from the dark bars (data not shown).

**Figure 3.**
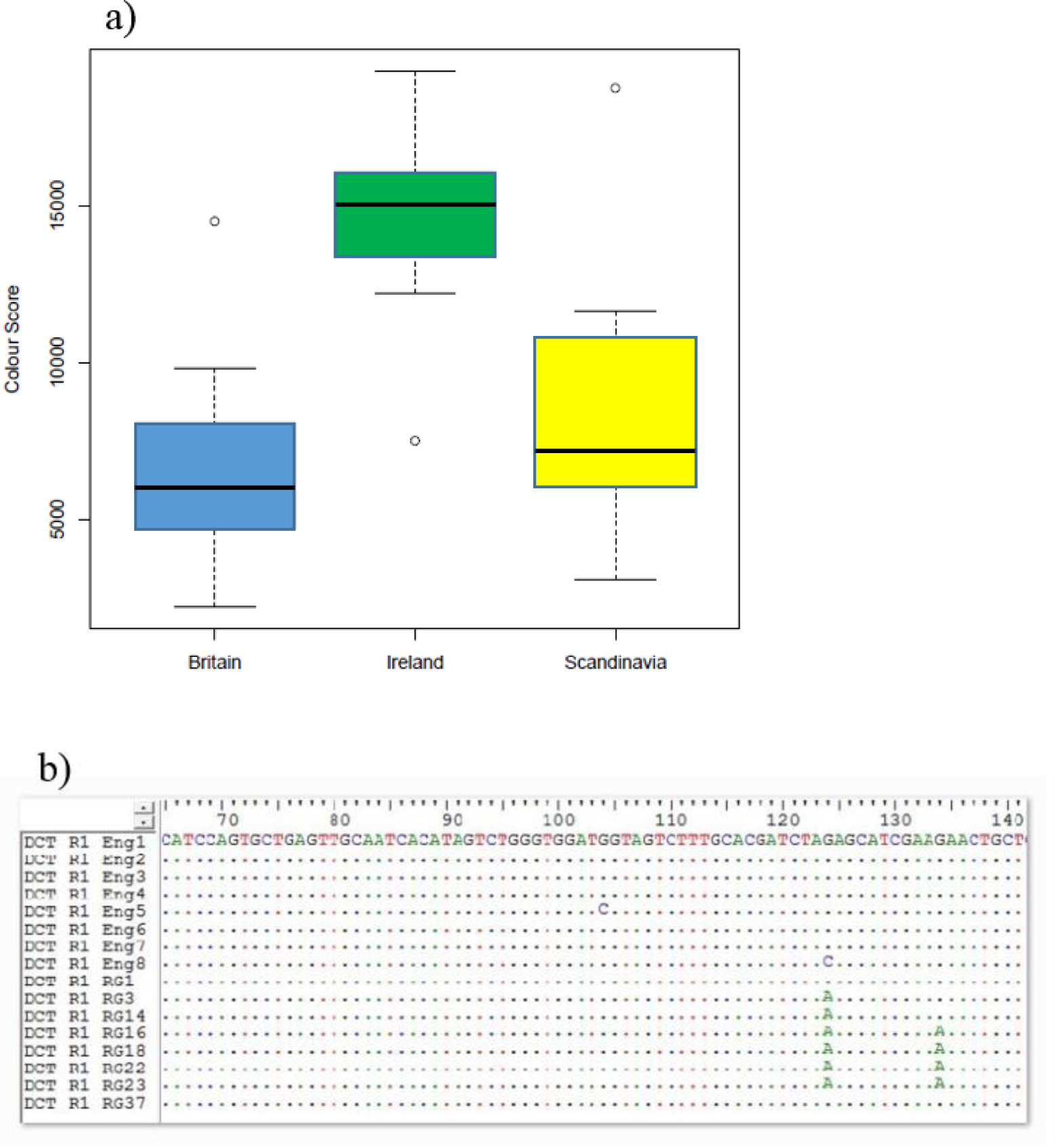
(**a**). Colour differences (photons reflected) on the light bars of the breast feathers among Scandinavian, Irish and British red grouse. A higher score means a lighter (less dark) feather. (**b**) Alignment of part of the coding nucleotide sequence of the DCT gene in British (sequence Eng1-8) and Irish (sequence RG1-through to 37) red grouse. A dot indicates perfect match with the top sequence. Note that the sequences are reversed in comparison to the chicken genome.

Four variable positions occurred in more than one haplotype (at positions 565, 568, 697 and 757) were detected in the coding sequence of the MC1R gene for a sample of Irish and British grouse (Table 2). All these substitutions constituted non-synonymous (amino acid changing) replacements. Furthermore, we found a difference among Irish and English birds in the coding sequence of the DCT gene (χ^2^=8.96, P = 0.003). Irish birds had an A (adenine) instead of a G (guanine) at position 124 in our alignment (in four cases this was also associated with the same substitution at position 134). When aligned to the homologous chicken sequence, the first substitution codes for a non-synonymous substitution conferring an amino acid replacement in Irish birds from Arginine to Lysine, the second substitution constituted a synonymous change (Fig. 3b).

**Table 2.** Nucleotide and amino acid replacements among British and Irish red grouse in the coding sequence of the MC1R gene. Numbers of sequences with the observed differences and the *P* value (Fisher’s exacts tests) for differences among Britain and Ireland. Position refers to nucleotide (amino acid in parenthesis) position from the start of the coding sequence.

| Position | Codon | Amino acid | Britain<br>(n=13) | Ireland<br>(n=15) | P value |
| --- | --- | --- | --- | --- | --- |
| **515 (189)** | CCC | Pro | 9 | 11 | 0.574 |
|  | AAC | Asn | 1 | 0 |  |
|  | ACC | Thr | 1 | 0 |  |
|  | TCC | Ser | 2 | 4 |  |
| **568 (190)** | GCT | Ala | 12 | 7 | 0.016 |
|  | ACT | Thr | 1 | 8 |  |
| **697 (233)** | TTT | Phe | 6 | 9 | 0.705 |
|  | CTT | Leu | 7 | 6 |  |
| **757 (253)** | GTT | Val | 2 | 3 | 0.099 |
|  | CTT | Leu | 11 | 12 |  |

### Gene ontology (GO) enrichment analysis

Gene ontology (GO) enrichment analysis was carried out with all 661 outliers. The gene ontology enrichment analysis results are shown in Table 3. There were no enrichment terms for Biological Processes. There were 12 enriched GO terms for Cellular Components. The most significantly enriched category was keratin filament.

**Table 3.** Gene ontology enrichment analysis for outlier gene identified using window-based F_ST_ outlier scans and pcadapt for Irish vs English red grouse.

| Enrichment FDR | Genes in list | Total genes | Functional Category/Pathway | GO term |
| --- | --- | --- | --- | --- |
| **Cellular Component** |  |  |  |  |
| 2.00E-08 | 10 | 16 | Keratin filament | GO:0045095 |
| 4.00E-07 | 29 | 206 | Intermediate filament | GO:0005882 |
| 4.00E-07 | 31 | 234 | Intermediate filament cytoskeleton | GO:0045111 |
| 2.80E-06 | 15 | 66 | Nucleosome | GO:0000786 |
| 1.10E-05 | 15 | 74 | DNA packaging complex | GO:0044815 |
| 9.30E-04 | 17 | 131 | Protein-DNA complex | GO:0032993 |
| 9.30E-04 | 39 | 482 | Polymeric cytoskeletal fibre | GO:0099513 |
| 6.40E-03 | 43 | 611 | Supramolecular complex | GO:0099080 |
| 6.40E-03 | 43 | 611 | Supramolecular polymer | GO:0099081 |
| 6.40E-03 | 43 | 610 | Supramolecular fibre | GO:0099080 |
| 2.20E-02 | 15 | 146 | Ribosomal subunit | GO:0044391 |
| 4.20E-02 | 16 | 173 | Ribosome | GO:0005840 |
| **Molecular Function** |  |  |  |  |
| 7.00E-06 | 54 | 604 | Structural molecule activity | GO:0005198 |
| 2.40E-02 | 16 | 138 | Structural constituent of ribosome | GO:0003735 |
| 4.30E-02 | 18 | 181 | Structural constituent of cytoskeleton | GO:0005200 |
| **KEGG pathway** |  |  |  |  |
| 3.50E-02 | 13 | 115 | Ribosome |  |

There were several genes including KRT18, KRT6A, KRT75, ENSGALG00000043689, KRT75L4, LOC112529929 and KRT8, which were enriched for seven GO enrichment categories including intermediate filament, keratin filament, intermediate filament cytoskeleton, polymeric cytoskeletal fibre, supramolecular complex, supramolecular polymer and supramolecular fibre. There were three categories that showed enrichment for Molecular Function which were Structural molecule activity, Structural constituent of ribosome and, Structural constituent of cytoskeleton. The KEGG pathway ribosome was also significantly enriched.

### Runs of Homozygosity

There were 7270 runs of homozygosity (ROH) detected. Of these 7112 were considered short/medium and 158 were considered long. Short/medium LROH were bounded between 100-1000 kb. There was a significant effect of population on LROH (p <0.001) and NROH (p=0.019) with a Tukey test revealing significant differences between the Recent vs Museum and English samples. There were 7112 Short/medium ROH, with an average (± sd) NROH of 309.2 (±163) per individual overall populations. The distribution of LROH per population is shown in Fig. 4 (a). The Recent samples show substantially longer short/medium LROH compared to the other populations. For short/medium ROH there was a significant effect of the population (p <0.001) on LROH following the same trend as for all ROH with Recent individuals being statistically different from English (p-adj <0.001) and Museum (p-adj <0.001) but English and Museum samples not being statistically different (p-adj=0.310) as shown by a Tukey test. Long ROH were above 1000 kb in length and ranged in size from 1002-6476 kb. Long ROH were recorded in 7 out of 9 English samples, 5 out of 8 Museum samples and 6 out of 6 Recent samples (Fig. 4 b). The SROH for long ROH segments against the frequency is shown in Fig. 4 (d) and indicated English individuals have the shortest and least frequent long ROH, followed by the Museum and then the Recent individuals. Whereas for short/medium ROH the Museum and Recent individuals were not differentiated in this regard, however, the English were still the lowest. There was no significant effect of population on long ROH length (p=0.076). The NROH was significantly affected by population (p <0.001) with a Tukey test revealing a significant difference between Recent and English. There is one individual as shown in Fig. 4 (d) that has more and longer ROH than all other individuals. This is an Irish individual from Co. Roscommon in the midlands and has 5 out of the 10 longest ROH. The Irish individual with the lowest ROH was from Co. Wicklow in the southeast.

**Figure 4.**
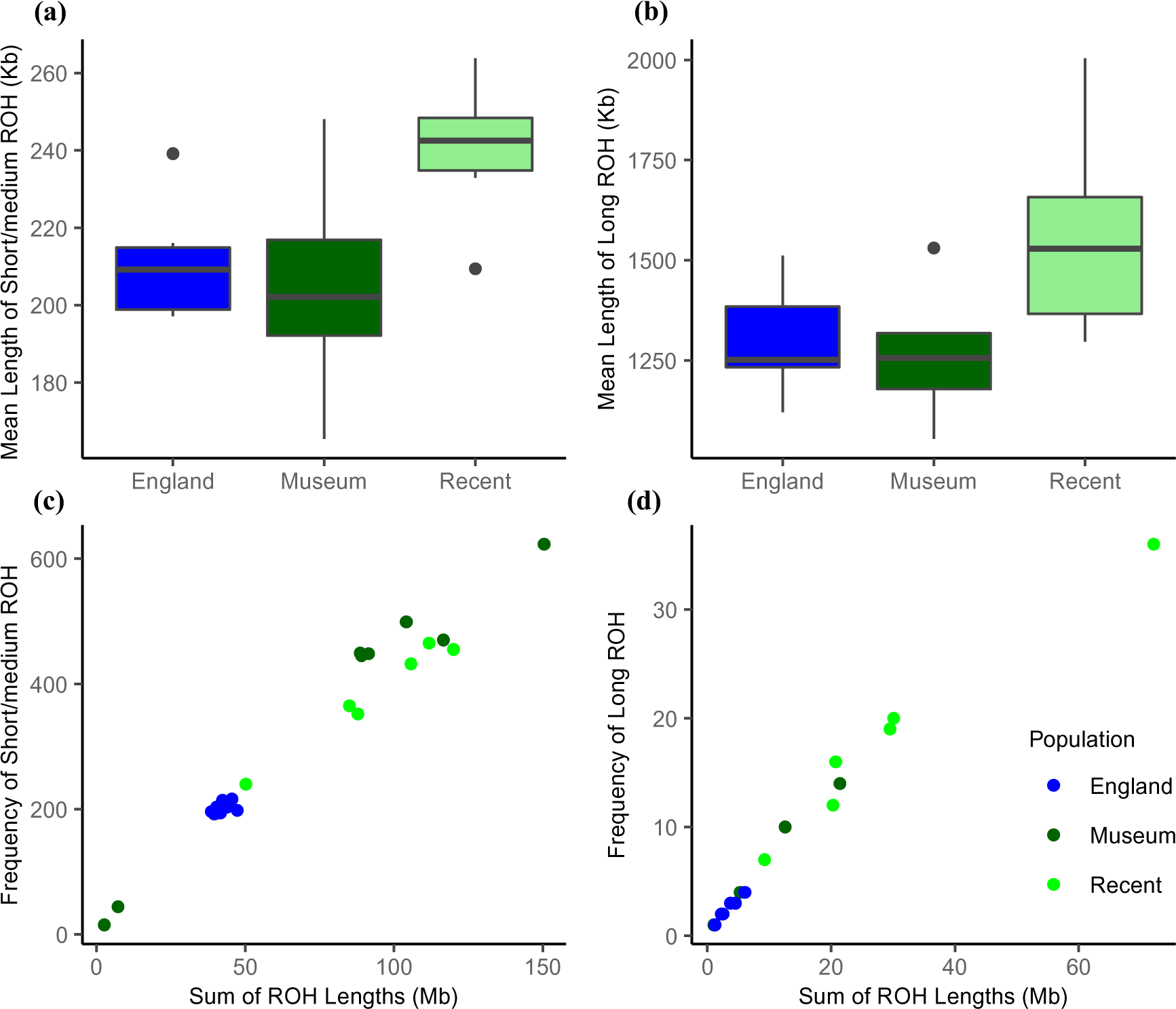
Boxplots showing the average length of (a) short ROH segments (100-1000 kb) and (b) long ROH segments (>1000 kb) for each population. Frequency of (c) short/medium and (d) long ROH against the sum of ROH for each individual with the colours indicating the different populations. The populations included were recent Irish, historic Irish and English.

## Discussion

Genomic analyses to assess the viability of species or population are becoming more accessible, however they still are not commonly used to design conservation programs (van Oosterhout, 2021). However, the application of genomics to conservation is growing (Hohenlohe et al., 2021). Key conservation data that can be gathered from population genomic studies include, genetic variation, effective population size, population structure, levels of inbreeding and demographic history (Hoban et al., 2022; Hohenlohe et al., 2021). Here we use that whole-genome data to illustrate some of these characteristics in a small and isolated populations and compare it to a large population. We then translate these results into practical conservation recommendations.

We show a clear population structure between Irish and British red grouse. More substructure was evident within the Irish population, due to the larger geographic extent of sampling and also likely due to different sampling times. With the Museum samples removed, there is more variation among English samples compared to Recent Irish (Fig S3(b)). The PCA showed two Irish individuals are noticeably closer to the English population relative to the other samples. In the ADMIXTURE (k=2) analysis, these Irish individuals showed the most admixture from the English population. We also showed using microsatellite variation that grouse from Northern Ireland were found in between birds from the Republic and birds from Scotland. These results indicate recent gene flow between the populations, which other studies did not find (Freeland et al., 2007; McMahon et al., 2012; Meyer-Lucht et al., 2016). Our results imply there has likely been human induced admixture from Britain in multiple Irish populations. If such translocations occurred, they would be non-sanctioned. The Species Action Plan for Red Grouse does note that they have occurred (National Red Grouse Steering Commitee, 2013).

Our genome-wide pairwise FST estimate between Recent Irish and English red grouse was 0.095 which is in line with previous studies of MHC-BLB genes(Meyer-Lucht et al., 2016) and somewhat higher than estimates for neutral microsatellite markers (McMahon et al., 2012) and neutral SNPs (Meyer-Lucht et al., 2016). Several peaks in the genome-wide window-based FST scans indicated selection as the dominant force shaping differentiation. In contrast to when genetic drift is the dominant force this plot would appear uniform. We found multiple peaks higher than the background level. Outlier analysis also identified 661 candidate genes under putative divergent selection. Several genes were found that occur very close together with particularly high ZFST indicating there could be an effect of linked selection.

Our analysis of runs of homozygosity (ROH) suggested, as expected, higher levels of inbreeding and lower levels of genetic diversity in the Irish samples, with long ROH indicative of lower genetic diversity. However, there was more spread in the Museum samples suggesting historic admixture has occurred in the Irish population. Within the Irish samples, there is one individual that shows relatively low SROH for long segments at 9238 kb. This is an admixed individual so this pattern is expected (Ceballos et al., 2018) and is from the Wicklow Irish population which has relatively high genetic diversity (McMahon et al., 2012). Another Irish individual had an extremely large long SROH (72147.95). This individual came from Roscommon, in the Irish midlands, which is an area that has experienced serious declines (Cummins et al., 2015).

We found evidence of divergence among Irish and British grouse in a number of genomic regions that harbour genes coding for ecologically important traits. At least 39 genes involved in immune response were identified as divergent among Ireland and Great Britain suggesting differential parasite mediated selection. These included B2M which is a component of class I MHC, involved in brain function (Smith et al., 2015), the presentation of peptide antigens to T-cells (Li et al., 2016) and is upregulated during viral infection (Yu et al., 2013). Previous research suggests MHC-BLB genes are locally adapted between the two populations (Meyer-Lucht et al., 2016). The gene RSFR, was identified and is likely to have bactericidal activity (Nitto et al., 2006). In chicken the protein encoded by this gene increases in macrophages following Salmonella infection (Sekelova et al., 2017). Another immune gene identified, FBXL8 is involved in tumour suppression in humans (Yoshida et al., 2021). There were four interleukins recorded, which are involved in the immune response. In addition, five chemokines and chemokine-like genes were identified. Chemokines are involved in the immune response and lymphocyte migration (Moser and Loetscher, 2001). One of these, CXCR4 was identified which is upregulated in red grouse with high T. tenuis burden (Webster et al., 2011).

We show here that Irish grouse are phenotypically paler than British red grouse and Scandinavian willow grouse. Among pigmentation genes identified as outliers, the MC1R gene, which encodes for the melanocortin 1 receptor is important for normal pigmentation in several organisms (Mundy, 2005). An interaction between this gene and the Agouti gene plays an integral role in pigmentation in many species (Harris et al., 2020). This gene is important in crypsis (Harris et al., 2020) and our sequencing of the coding part showed it to be differentiated between Irish and British red grouse. Another candidate gene in the melanin pathway DCT also showed divergence when the coding region was sequenced. HSDL1 was identified as an outlier and may play a role in pigmentation as it is involved in the maintenance of secondary sexual characteristics and sex differentiation by metabolising hormones (Gloux et al., 2019). The genes HPS1 and SNAPIN were also extracted as outliers and in humans are involved in melanin production and melanosome biogenesis (Martina et al., 2003), (Setty et al., 2007). Mutations in HPS1 have been shown to cause changes in mice pigmentation (Nguyen and Wei, 2007).

Heather is the primary food source of red grouse in Scotland (Savory, 2008) and Ireland (Lance and Mahon, 1975). It is believed there is a lower availability of heather in Ireland compared to Britain. With this in mind, several genes involved in food intake and energy control were identified. The expression of GALR2 and GALR3 which were identified as outliers, in tissues involved in the maintenance of glucose homeostasis is regulated by short-term food deprivation (Kołodziejski et al., 2021). Also, the gene GHRL is significantly associated with body weight, body weight gain, and feed conversion ratio in chickens (Jin et al., 2014). HCRT is potentially involved in energy balance in chickens (Miranda et al., 2013). This gene was identified using both methods. The gene DRD5 may be involved with feeding behaviour in chickens (Jiang et al., 2021). Lastly, the gene RHNO1 is downregulated in chickens which are very efficient with feed intake (Yang et al., 2020). Other genes potentially involved in energy control and food intake include NPW (Bu et al., 2016; Nagata-Kuroiwa et al., 2011; Takenoya et al., 2010), GRIN1 (Liu et al., 2012) and MLST8 (Jacinto et al., 2004).

The gene ontology enrichment analysis showed divergent genes were significantly enriched for GO categories mostly concerning Cellular Components. The most enriched categories involved the keratins and included intermediate filament. Of the genes which were enriched, 25 were keratins. Intermediate filaments are essential for maintaining cell structure and plasticity (Herrmann et al., 2009). In birds, they contribute to the formations of beaks, feathers, claws and scales (Ng et al., 2014). Bird feathers consist of α and β keratins, with β keratins unique to birds and reptiles (Wu et al., 2015) There was one β-keratin-like gene LOC431320 that was enriched for six categories and one β-keratin related protein BKJ which was enriched for six also. There were also five unnamed feather keratin-like genes which showed significant enrichment for several categories. This indicates that red grouse in Ireland and England are differentiated according to functions involving keratin suggesting a role in feather formation.

Two recent studies on red grouse (McMahon et al., 2012; Meyer-Lucht et al., 2016) concluded that the Irish and the British populations of red grouse should be treated as separate “Management Unit” (MU), as they unravelled significant and important differentiation among these two populations (Moritz, 1994). This study agrees with this and has added to the evidence on important differentiation with a list of candidate genes under putative divergent selection.

Unlike other studies, this study shows recent gene flow from Britain to Ireland especially for the north of Ireland and substantial admixture in at least two of the southern populations. Regardless, selection seems to be playing an important role in shaping the differentiation between these two populations. The two populations show divergence in genes with important biological functions including the immune response, pigmentation and food intake. This suggests red grouse in Ireland are adapted to the local conditions of potentially divergent parasite communities as well as different levels of heather in each habitat. The introduction of maladapted phenotypes and genotypes can result in outbreeding depression. This is when admixed individuals have reduced fitness (Frankham et al., 2011) and this could exacerbate conservation efforts. In addition, the contemporary Irish population has significantly more and longer ROH than the English individuals. The ROH analysis indicated higher inbreeding in Ireland, that this has potentially increasing over time and that there is likely reduced genetic diversity in the Irish population (Freeland et al., 2007).

It is clear that for the Irish population with increased levels of inbreeding, reduced genetic diversity and continuing populations declines (Cummins et al., 2015) conservation action is needed (National Red Grouse Steering Commitee, 2013). There is still a large enough population of red grouse in Ireland so that habitat management alone could be sufficient to increase the species but this will not be the case if there are further declines. The estimate from 2016 of an Ne of 100 means that if populations continue declining it may not be possible to conserve the Irish red grouse due to loss of genetic variation (Lynch and Lande, 1998; Meyer-Lucht et al., 2016). It is therefore critical that this does not happen.

It is advised here, in agreement with previous studies (McMahon et al., 2012; Meyer-Lucht et al., 2016) that translocations from Britain be avoided due to the divergence in several important genes and the risk of introducing genetic variation from maladapted birds. This recommendation is also in line with what is stated in the Species Action Plan (SAP), and IUCN guidelines for the ‘Guidelines for Reintroductions and Other Conservation Translocations’ (IUCN/SSC, 2013). While these results indicate that translocations should not occur in order to preserve the divergent adaptive variation, genome-wide variation in populations should also be considered(Kardos et al., 2021). To preserve genetic variation, gene flow between the small and fragmented Irish populations should be restored and at the bare minimum retained through habitat management. Restoring gene flow between the four main populations in Ireland through translocations could help minimise the effects of inbreeding. If this occurs, managers must ensure that the chosen source population is healthy enough to support the removal of individuals (National Red Grouse Steering Commitee, 2013). It should also be ensured that maladapted genotypes are not moved around Ireland to isolated populations.

These data indicate that there has been recent inbreeding in the contemporary Irish population. There is evidence of recent gene flow from the English population into some of the Irish individuals and this may cause negative effects through outbreeding depression. Our study demonstrates the applications of WGS with its use for taxonomy and the conservation of an endangered species.

## Materials and Methods

### Samples preparation

#### Study samples

Whole genome sequences of 23 individuals were analysed in this study. Details of these are shown in Table S1 and Fig. 1. Eight individuals were sampled from museum skins at the Natural History Museum, Dublin. The museum samples were from Irish individuals and were all catalogued between 1881-1882. There were also feather samples from four Irish birds which were collected in 2007. Two more samples, one feather and one muscle were obtained from Irish birds in 2020. In addition, there were nine sequenced genomes from English red grouse from a previous study (Kozma et al., 2019) which were from two spleen and seven liver samples. These samples were collected from the Yorkshire Dales in the UK in 2013. In this report, the samples will be referred to as English, Museum and Recent. English referring to the English samples, Recent referring to the feather and muscle samples collected in Ireland since 2007 and Museum referring to the Irish samples originally catalogued between 1881-1882. The Recent and Museum samples are referred to collectively as the Irish samples. For analyses of colour scores and sequencing of pigmentation gene candidate loci see below under “pigmentation genes”.

#### DNA extractions

The DNA extraction protocol varied for the different samples. For the 9 English samples, DNA extractions were carried out as described in Kozma *et al*. 2019 with a Qiagen DNeasy Blood & Tissue Kit® following the manufacturer’s instructions (Qiagen). The quality of DNA was assessed on an agarose gel and a Quibit® Fluorometer. Extractions for the recent muscle samples were carried out using a Thermo Scientific KingFisher Cell and Tissue DNA Kit following the manufacturer’s instructions for the KingFisher Duo purification system. Before commencing the tissue samples were broken down mechanically using a scalpel. DNA yield was assessed using a Qbit 3.0 Fluorometer. DNA extractions for the feather and historical toepad samples were carried out using a QIAamp DNA Micro Kit following the manufacturer’s protocol for the isolation of genomic DNA from tissues less than 10mg with some deviations. During the lysate stage 20ul of DTT was added to aid in tissue breakdown. There was an extra incubation step for 10 minutes at 72°C DNA during the lysis stage. Lastly, the final dilution step was split into two with 30ul added each time and a centrifuge in between.

#### Library preparations

For the nine English samples, library preparation was carried out using the Illumina TruSeq protocol as described in Kozma *et al*. 2019. The muscle and feather sample libraries were prepared with the Illumina TruSeq PCR-free protocol. Library preparations for the museum samples followed the protocol “Illumina Sequencing Library Preparation for Highly Multiplexed Target Capture and Sequencing” (Meyer and Kircher, 2010) which was modified by adding USER enzyme and dual indexing (Briggs et al., 2010). For detailed descriptions of all laboratory procedures for degraded DNA from historical samples, see Irestedt *et al*. (2022) (Irestedt et al., 2022).

#### Sequencing

The English samples were sequenced using an Illumina HiSeq 2500. They were pair ended reads of 125bp with a target insert size of 350 bp (Kozma et al., 2019). The feather and tissue samples were sequenced on a quarter lane of S4 NovaSeq6000 (NovaSeq Control Software 1.7.0/RTA v3.4.4) using the standard protocol. They were pair ended with a read length of 2×150bp. The museum samples were sequenced on one NovaSeq6000 (NovaSeq Control Software 1.7.0/RTA v3.4.4) and were also pair ended.

#### Microsatellites

Allelic variation at 15 neutral microsatellite loci were scored for 7 birds from Scotland (Aberdeenshire) and 15 birds sampled in Northern Ireland. The loci scored and the methods used were the same as described in McMahon et al. (2012) (McMahon et al., 2012) and Johansson et al. (2012) (Johansson et al., 2012). Briefly feathers were collected in the field either as pick-ups of moulted feathers or from shot birds. Feathers were transported to the lab were DNA was extracted from the feather roots using a salting out protocol (Johansson et al., 2012). Allelic variation at the following loci were scored as previously described: ADL142, ADL230, ADL184, BG15, BG16, BG 18, TUT 1, TUT2, TUT3, LLSD2, LLSD3, LLSD4, LLSD6, LLSD7 and LLSD8 (Johansson et al., 2012; McMahon et al., 2012).

#### Pigmentation genes

Following the methods by Skoglund and Höglund (2010)(Skoglund and Höglund, 2010) we obtained sequence data from two genes involved in the melanogenesis pathway: Melanocortin 1 Receptor (MC1R) and Dopachrome tautomerase (DCT). We extracted genomic DNA from feathers from 16 birds from Teesdale, Yorkshire, UK and 25 birds from Ireland and used the primers described in Skoglund and Höglund (2010) to obtain PCR-products of the desired length. For MC1R we used the primer pair MSHR80-MSHR9 and for DCT we used DCTF2-DCTR1. Polymerase chain reactions (PCR) were run as in Skoglund and Höglund (2010) and PCR-products were Sanger sequenced on a MegaBace 1000 capillary instrument (GE Healthcare, Wakeusha, WI, USA) using Dyenamic ET terminators (GE Healthcare) according to the manufacturer’s recommendations.

#### Mapping and SNP calling

Mapping and SNP calling was carried using a bioinformatic pipeline which followed the suggested best practices workflow by the *Genome Analysis Toolkit* (*GATK*) (Van der Auwera *et al*. 2013, BroadInstitute 2020).

Sequencing output fastqc files were merged for each individual. Quality checks were then carried out on these samples using the tool *Fastqc* (v0.11.9) (Andrews, 2010) which provides an overview of the data quality and highlights potential issues. The software *Trimmomatic* (v0.36) (Bolger et al., 2014) was used to identify and remove adapter sequences as well as to remove low-quality areas. *Trimmomatic* was used with the following settings: [LEADING:5 TRAILING:5 SLIDINGWINDOW:4:15 MINLEN:50]. This resulted in a window 4 base pairs wide being read and subsequently cut when average quality dropped below 5 and then reads below 50 base pairs in length were also dropped. This resulted in four files per individual consisting of paired forward and reverse reads and unpaired forward and reverse reads. For further analysis, the paired read files were used.

The resulting paired forward and reverse reads were then aligned to the reference chicken genome using the *Burrows-Wheeler Aligner* (*BWA*) (v0.7.17). *BWA* was ran using the mem algorithm with long-read support for split alignment on 20 threads [-t] and Picard compatibility [-M]. The output is one BAM file for each individual. The *MarkDuplicates* was then used to identify and mark the duplicates so that they would be ignored in later steps. The tool *index* from SAMtools (v.1.5) (Li et al., 2016) was used to index the resulting files. *QualiMap* (v.2.2.1) (García-Alcalde et al., 2012) was used to assess read quality.

Variant calling was carried out according to the steps outlined by GATK (v4.2.0.0) (Van der Auwera et al., 2013) using three tools – *HaplotypeCaller, CombineGVCFs* and *GenotypeGVCFs. HaplotypeCaller* was used on each sample with [-ERC GVCF] to create an intermediate g.vcf to be run through *GenotypeGVCFs.* The output of *HaplotypeCaller* is a g.vcf for each chromosome per individual. Before these can be genotyped they must be combined into one file with *CombineGVCFs*. The file is then joint genotyped with *GenotypeGVCFs* resulting in one single variant call format (VCF) with all samples.

The next step was filtering the vcf. Several tools were used for this. SNPs were extracted using *SelectVariants* in two steps. Firstly, [--select-type-to-include SNP] was used to select SNPs and [--restrict-sites-to BIALLELIC] to only include biallelic sites, excluding multiallelic sites and these filtered sites were excluded [--exclude-filtered true]. Then *VariantFiltration* was used with the filtering criteria: [QUAL < 30, MQ < 40.00, SOR > 4.000, QD < 2.00, FS > 60.000, MQRankSum < −12.5, ReadPosRankSum < −8.000]. Finally, SNPs in repetitive regions were removed with *BEDTools* (v.2.27.1). For this, the chicken repeat track was used. This could potentially result in some error as it may not correspond exactly to the repetitive regions in red grouse. For this the command *subtract* was run with default settings and the chicken repeat track as the reference. A similar process was carried out for indels where *SelectVariants* was used to select the indels [--select-type-to-include INDEL] and filtered sites were excluded [--exclude-filtered true]. *VariantFiltration* was then ran with the criteria: [QUAL < 0, MQ < 40.00, SOR > 10.000, QD < 2.00, FS > 200.000, ReadPosRankSum < −20.000]. Then BEDTools (v2.27.1) (Quinlan and Hall, 2010) was used to remove SNPs within 5bp of an indel using the commands *window* and *subtract.* The flag [-w 5] was used with *window* to search for overlapping windows between the output SNP file and the output indel file from the previous steps. It marks 5 base pairs upstream and downstream of each overlapping feature in the SNP file which were then removed. Using *subtract* the output file is then written to a vcf which has been hard filtered, with SNPs in complex repetitive regions removed as well as SNPs within 5bp of indels.

The last filtering step is carried out using *VCFtools* (v0.1.15) (Danecek et al., 2011). The Z and W sex chromosomes were removed. The file was filtered to include sites with a minor allele frequency greater than or equal to 0.5 [--maf 0.05] and sites, where the max missing data for one locus was more than 10%, were cut [--max-missing 0.9]. This dataset was now ready for analysis. The same dataset was then filtered to remove linkage disequilibrium at a threshold of 0.4 using *PLINK* (v1.9) (Purcell et al., 2007).

### Analyses

For some analyses, the museum samples are omitted for two reasons. Firstly, the levels of deamination in the museum samples were not quantified or filtered for. In historic samples, there can be damage at CpG regions due to cytosine deamination and conversion into uracil (van der Valk et al., 2019). This could lead to incorrect conclusions regarding divergence. Also since there was a c. 120-year time gap between the Irish samples this could have affected the results. Therefore in some analyses where it is considered this could have a large effect the museum samples were left out.

#### Genetic diversity

Nucleotide diversity (π) was calculated among the English and Irish samples. For this, the Irish samples were split into Recent and Museum samples to investigate whether there had been a drop in genetic diversity over time in Irish birds. This was carried out in *VCFtools* (v.0.1.15) (Danecek et al., 2011) with the command [--window-pi 15000] to measure nucleotide diversity per 15 kb window.

#### Population structure and local adaptation using pcadapt

To investigate population structure and identify evidence of local adaptation a package called pcadapt (Privé et al., 2020) was implemented in R (v.4.0.4). Firstly, a Principal Component Analysis (PCA) with all samples was carried out to visualise genetic variation. Following this, a second PCA was carried out with only the Recent Irish and English samples. The software pcadapt then detects outliers based on the principal components (PCs). It computes *k* PCs based on the genetic variation which explain population structure. Initially, 10 PCs were used and this was graphically inspected using a scree plot (Fig. S2(a)) to assess the variance explained by each PC and also with a score plot to so see how individuals clustered. It was decided to retain two PCs which were believed to be related to population structure. SNPs were then regressed against the two PCs and z-scores were calculated. Mahalanobis’ distances of these z-scores were then calculated and converted into p-values. The p-values for each SNP were transformed into q-values with the R package *q-value* and a false-discovery rate (FDR) of 0.05 was used to detect outliers. For the outlier analysis, only the Recent Irish samples and English samples were used which has been pruned for linkage disequilibrium.

#### Analysis of Admixture

To estimate ancestry, the software ADMIXTURE (Pritchard et al., 2000) was used. For this analysis, the BED file was used that had been pruned for linkage disequilibrium which is the requirement for this software. This was run for populations (k) set to 1-5. By using [--cv] a cross-validation error for each k was obtained. This error estimate gives the accuracy of the estimate with lower values being more accurate. The k with the lowest value is considered to be the most likely number of populations. The ADMIXTURE results were then visualised in R (v.4.0.4) using *tidyverse* and *ggplot2*.

#### Fst outlier analysis

To assess genetic differentiation further, pairwise fixation indices (F_ST_) (Weir and Cockerham, 1984) were calculated for Recent Irish vs English red grouse. F_ST_ is a measure of population differentiation ranging from 0-1, with 0 being the most similar and 1 being the most differentiated. This was calculated using *PLINK* (v1.9) (Purcell et al., 2007) and the [--fst]function. In addition, window-based F_ST_ across the genome was calculated for 15 kb windows using also with the Recent Irish and English samples. This was done with *vcftools* and the options[--weir-fst-pop] and [--fst-window-size 15000]. The dataset had not been thinned for linkage disequilibrium. The F_ST_ scores were transformed into ZF_ST_ scores. These show how many standard deviations a data point is from the mean. The results were then visualised in R with the *qqman* package.

These window-based ZF_ST_ values were used to determine genes in regions with high F_ST_. Windows with ZF_ST_ >3 were considered outliers and were mapped against the chicken genome. Genes within 5 kb above and below these windows were extracted. Particular attention was paid to genes with a function in immune response or plumage colour functions as these have been previously studied in the species and put forward as being under divergent selection (Meyer-Lucht et al., 2016; Skoglund and Höglund, 2010)

#### Phenotypic colour quantification

We collected abdominal feathers from the lower part of the neck from seven birds from Scandinavia (three from Kongsvoll, Norway, three from the adjacent islands Frøya and Hitra, Norway and one from Transtrand, Sweden), eight birds from Ireland (three from Co. Wicklow, one from Co. Limerick, and five birds from Northern Ireland) and 20 birds from Teesdale, Yorkshire, UK and took 2-8 reflectance measurements per bird on the light and dark bars of each feather, respectively. Reflectance measurements were obtained with an Ocean Optics S2000 diode array spectrophotometer from surfaces illuminated by a TOP Sensor DH-2000 combined deuterium-halogen light source according to the procedures described in(Håstad and Ödeen, 2008) with the exception we limited our measurements to only be taken at a 90° angle to the feather surface. Reflectance data (roughly number of photons emitted while positioning a c. 1 mm^2^ reflectance probe on the feather) were averaged per bird from two kinds measurements of each contour feather (i.e. light and dark bars on each feather). Since we were mainly interested in comparing potential colour differences among the three populations (Scandinavia, Ireland, Great Britain) we report reflectance as average number of photons emitted per measurement rather than full reflectance spectra.

#### Colouration

We tested for differences among regions using Kruskall Wallis one-way analysis of variance and used Wilcoxon-tests for pair-wise comparisons.

#### Sequence data

Sequences were assembled and manually edited with BioEdit (http://www.mbio.ncsu.edu/BioEdit/bioedit.html) and sequence alignments were created with Clustal V using default parameters. Reading frame was inferred by comparing with the chicken (*Gallus gallus*) genome sequence (International Chicken Genome Sequencing Consortium 2004) and all alignments were manually inspected for non-synonymous and synonymous polymorphisms that clustered within either population. Haplotypes were inferred using PHASE (Sheet and Stephens 2006) with 1000 iterations and a 100 generation burn-in. We tested for unequal distribution of individual SNPs by Fisher Exact tests.

#### Gene ontology (GO) enrichment analysis

ShinyGo v.0.61 (Ge et al., 2020) was used to carry out gene ontology (GO) enrichment analysis based on a false discovery rate (FDR) of <0.05 as statistical significance. This analysis was run to identify gene ontology enrichment in Biological Processes, Molecular Function, Cellular Components and KEGG pathways.

#### Runs of Homozygosity analysis

To detect runs of homozygosity (ROH) there were two different criteria used. This was done with *PLINK* and the [--homozyg] option. The scanning window was 50 SNPs wide and set so that SNPs that were over 1000 kb apart could not be considered in the same ROH. Only stretches containing at least 50 SNPs that were 100 kb were considered. Following this, the ROH output was split into short/medium and long ROH. Short/medium ROH was defined as homozygous stretches between 100-1000 kb. These are signs of past demographic effects such as ancient inbreeding and bottlenecks whereas long ROH are considered signs of recent inbreeding (Ceballos et al., 2018). Long stretches of ROH were classed as those greater than 1000 kb. The total number and average length of ROH per individuals and total length were visualised in R using the packages *ggplot2* and *tidyverse*.

There may be an issue with heterozygote sites being incorporated into the sequencing data as an effect of post-mortem DNA damage, which would break up ROH. We have taken precautionary steps to prevent this by USER-treating during library preparation and have quite low degrees of DNA damage indicated in mapdamage2 (Jónsson et al., 2013). However, it may still introduce false positives in our museum samples, which would cause shorter ROH.

## Acknowledgments

Historic samples were generously provided by The National Museum of Ireland – Natural History in Dublin with the help of Dr Aidan O’Hanlon and Paolo Viscardi. Fresh samples were provided by Séamus Butler and Frank Reynolds. Feather samples were in storage at Uppsala University. The English samples were produced from a previous study and were sequenced at the SNP&SEQ technology platform at Uppsala University (Kozma et al., 2019). Library preparations for the museum samples occurred at the Department of Bioinformatics and Genetics at the Swedish Museum of Natural History in Stockholm, Sweden. Library preparations for the muscle and feather samples were carried out by NGI Solna. The authors acknowledge support from the National *Genomics* Infrastructure in Stockholm funded by Science for Life Laboratory, the Knut and Alice Wallenberg Foundation and the Swedish Research Council, and SNIC/Uppsala Multidisciplinary Center for Advanced Computational Science for assistance with massively parallel sequencing and access to the UPPMAX computational infrastructure.

## Notes

https://www.ncbi.nlm.nih.gov/bioproject/?term=PRJNA512999

https://dataview.ncbi.nlm.nih.gov/object/PRJNA991980?reviewer=npbkhs092n69c375qqi78bast5

https://doi.org/10.5061/dryad.jm63xsjgk

https://datadryad.org/stash/share/WE9rIoiFOmNIBCd0Q6aLQrr4U5Eg-k8VhSQAg9GpILQ

### Competing Interest Statement

The authors have declared no competing interest.

