## Supplementary Information for "Inbreeding or outbreeding depression? How to manage an endangered and locally adapted population of red grouse *Lagopus scotica*"

**Supplementary Material**

**Results: Outlier Analysis**

*Immune response*

There were up to 39 genes that were potentially involved in the immune response. The two genes which showed the highest differentiation were ADAM8 (ADAM metallopeptidase domain 8) with a ZFST of 9.52 and RSFR (leukocyte ribonuclease A-2) with a ZFST of 9.2. The gene FBXL8 was the only immune response associated gene found by both methods and is associated with cancer in mammals. Of these genes, 11 were associated with microbial or viral infection chickens and 19 with the response to pathogens in mammals. At least 5 were associated with cancer, four in mammals and one in chicken.

*Pigmentation*

Regarding pigmentation, several genes were identified and are shown in Tables S3 and S4 (highlighted in orange). Those with the highest ZFST of 10.26-8.25 were LIPML1, LIPML2 and LIPML3 (lipase member M-like 1,2,3). These are potentially involved in the final steps of keratinocyte differentiation. Also HSDL1 (hydroxysteroid dehydrogenase-like 1) which is involved in the maintenance of secondary sexual characteristics, which in birds includes plumage colour was identified as an outlier by both outlier methods (Table S2). There were 5 feather keratin-like genes identified. Of these three (KRT75L4, KRT6A, KRT75) were identified in both methods. MC1R which codes for melanocortin 1 receptor was identified by pcadapt. This is a well-known and studied gene involved in pigmentation. When taking a closer look at the region containing this gene it was found that there was a SNP within the MC1R region with a particularly high FST of 0.3 indicating high differentiation for one SNP in this gene. Other notable genes involved in melanin production include HPS1 and SNAPIN.

*General genes and outlier method overlaps*

There were also many other genes identified with other important functions (highlighted red in Tables S3 and S4). There were 8 genes associated with the eye, most often process in the retina in both chickens and mammals. There were also some genes involved in male fertility and spermatogenesis in humans and/or mice and also one gene associated with fertilization. There were genes involved in stress responses including hypoxia in chickens and cold stress. Lastly, there were seven genes involved in various behavioural traits such as food intake, sleep and memory and energy balance. Of these, genes with particularly high differentiation include RDH8, DDX25 and DNAH3.

There were also a further 9 genes that were found by both methods and did not fall into any of the above categories. These are shown in Table S2. There was one keratin-like gene and two histones. Also PIGC, HS6ST3, HEBP1 and S100B.

**Discussion: Outlier Analysis**

Other *interesting* functions

Genes with other important functions were involved in reproduction, processes in the eye and food intake. Previously, it was found that the EDIL3 gene was under divergent selection between grouse species (1). This gene was not identified here. However, the gene is involved in egg mineralization in chickens along with another gene MFGE8 which was identified here as an outlier (2).Interestingly, several genes involved in the eye were also detected. SIX5 is expressed in adult human eyes (3). ATOH7 is involved in retina development in chickens (4)and FOXG1 is in mice (5), both were identified as outliers. HPS1 plays a role in the iris and has underwent intensified evolution in the development of barn owl eyes (6). RLBP1 is required for the correct functioning of rod and cone photoreceptors in humans (7) and is very important for vision in mice (8). Lastly, PDC is present in mammal retinas and is associated with vision in mice and humans (9, 10). There were also two genes associated with high altitude one of which is the main candidate gene for hypoxia in Tibetan chicken - FOXG1 (11). The second is EIF2AK1 and is potentially involved in the adaptation to hypoxic stress in Tibetan chickens (12). There were several genes involved in spermatogenesis and sperm function in mice such as DDX25 (13) and SYT6(14). Also, there were two genes involved in spermatogenesis in humans TSGA10 (15) and TMEM203 (16).

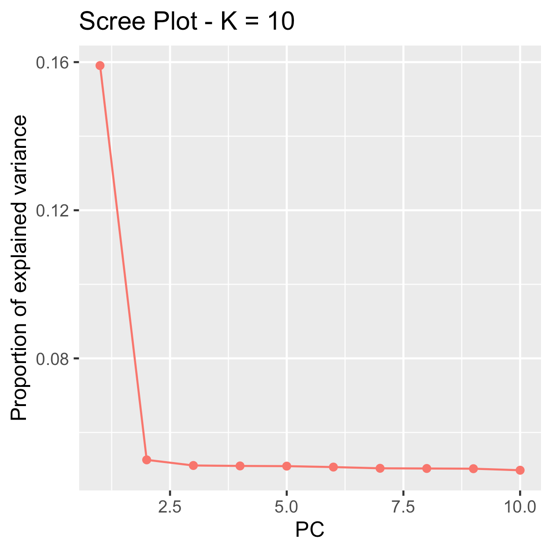

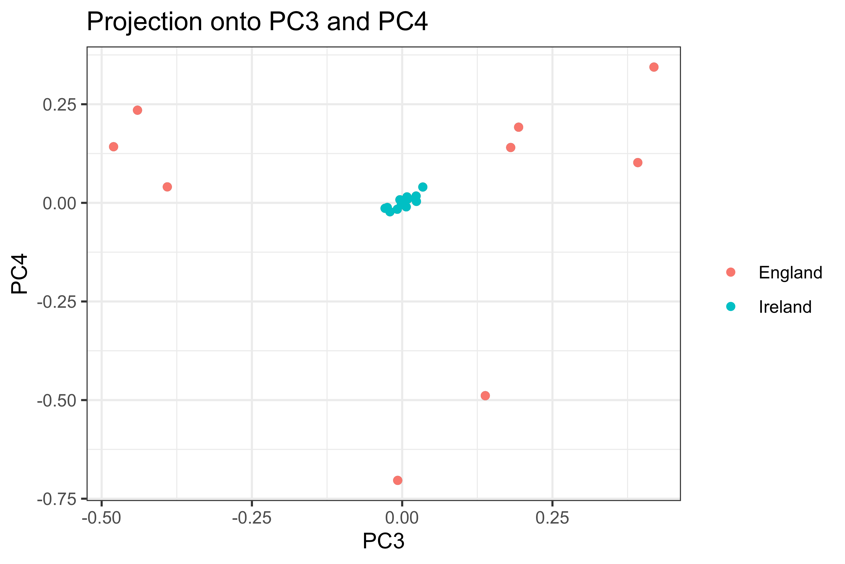

(b)

(a)

**Figure S1.** **(a)** Scree plot showing the variance explained by the first 10 PCs including all samples (Recent, Museum and English) using pcadapt. **(b)** PCA score plot showing PC3 and PC4 for English and all Irish samples.

(a)

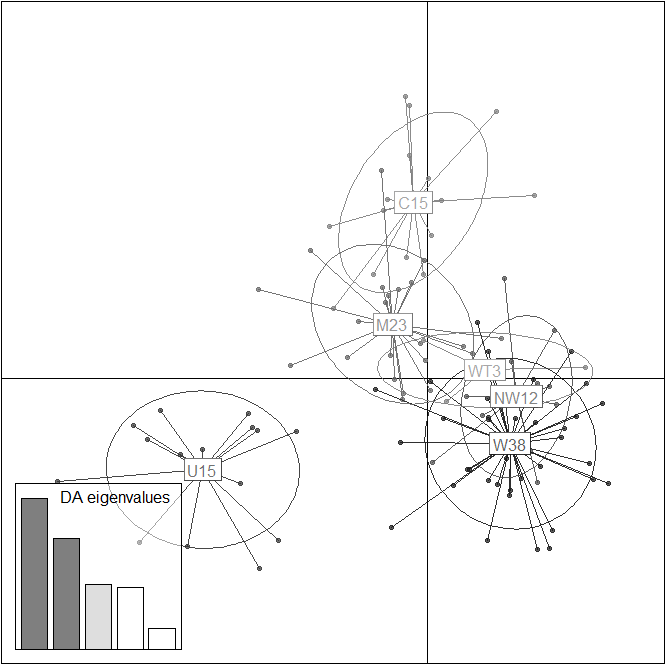

(b)

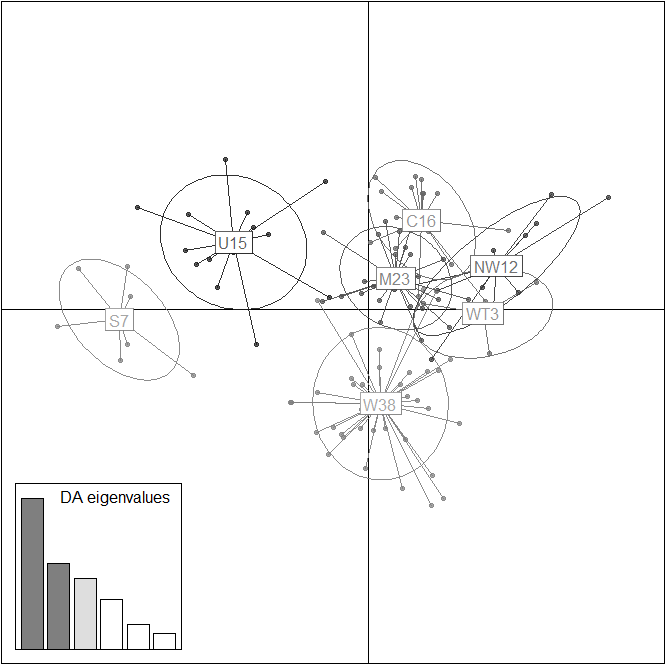

**Fig. S2.** PCA-plot of microsatellite allelic variation among Irish red grouse populations (a) and when adding seven samples of birds from Scotland (b). Population codes: C- Cork, M- Munster, NW- North West, U-Northern Ireland, W- Wicklow, WT-West, S-Scotland.

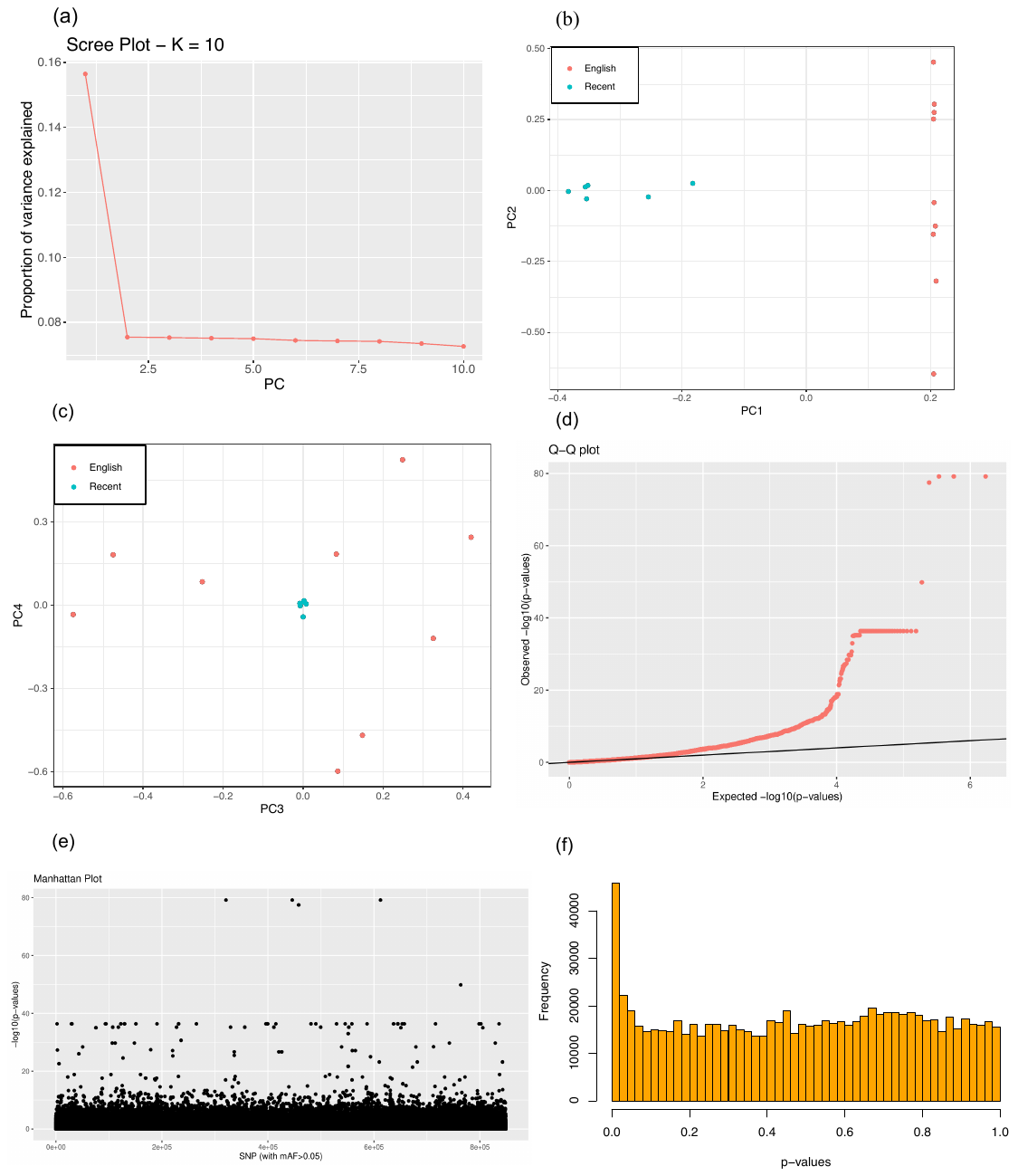
**Figure S3.** **(a)** Scree plot showing the variance explained by the first 10 PCs for the comparison of contemporary Irish and English samples. Score plots for **(b)** PC1 and PC2 and **(c)** PC3 and PC4 and **(d)** qq plot of -log10(p-values) for the SNPs, **(e)** Manhattan plot of -log10(p-values) and **(f)** histogram of the frequency of p-values.

**Table S1** Information on samples.

| **Sample ID** | **Museum ID** | **Location** | **Sample type** | **Sample year*** | **Mean coverage (x)** |
| --- | --- | --- | --- | --- | --- |
| IreScot1 | - | Clare | Feather | 2006/2007 | 21.0 |
| IreScot2 | - | Monaghan | Feather | 2007 | 22.4 |
| IreScot3 | - | Cork | Feather | 2007 | 21.9 |
| IreScot5 | - | Roscommon | Feather | 2007 | 23.3 |
| IreScot6 | - | Knockmealdowns, Tipperary | Feather | 2020 | 21.1 |
| IreScot9 | - | Wicklow | Muscle | 2020 | 20.9 |
| MusScot1 | 1881.590.1 | Sligo | Toepad | 1881 | 23.7 |
| MusScot2 | 2003.30.9 | Tipperary | Toepad | 1881 | 23.7 |
| MusScot3 | 1881.589.1 | Offaly | Toepad | 1882 | 29.0 |
| MusScot4 | 2003.30.17 | Monaghan | Toepad | 1881 | 23.6 |
| MusScot5 | 1881.594 | Kerry | Toepad | 1881 | 23.8 |
| MusScot6 | 2003.30.27 | Wicklow | Toepad | 1881 | 23.7 |
| MusScot7 | 2003.30.23 | Limerick | Toepad | c. 1881 | 24.0 |
| MusScot9 | 2003.30.24 | Knockmealdowns, Waterford | Toepad | 1881 | 19.7 |
| Scot1 | - | Feetham, Yorkshire Dales, UK | Liver | 2013 | 26.5 |
| Scot2 | - | Feetham, Yorkshire Dales, UK | Liver | 2013 | 26.9 |
| Scot3 | - | Feetham, Yorkshire Dales, UK | Liver | 2013 | 30.3 |
| Scot4 | - | Feetham, Yorkshire Dales, UK | Liver | 2013 | 26.3 |
| Scot5 | - | Gunnerside, Yorkshire Dales, UK | Liver | 2013 | 29.6 |
| Scot6 | - | Gunnerside, Yorkshire Dales, UK | Spleen | 2013 | 26.8 |
| Scot7 | - | Gunnerside, Yorkshire Dales, UK | Spleen | 2013 | 27.2 |
| Scot8 | - | Gunnerside, Yorkshire Dales, UK | Liver | 2013 | 27.4 |
| Scot9 | - | Gunnerside, Yorkshire Dales, UK | Liver | 2013 | 28.6 |

**Table S2**. Outlier genes that were recorded by both outlier methods.

| **Chrom-osome** | **ZF_ST_** | **Ensembl Gene ID** | **Gene name** | **Description** | **Source** |
| --- | --- | --- | --- | --- | --- |
| 10 | 7.4 | ENSGALG00000002097 | LOC415324 | epididymal protein-like | - |
| 27 | 6.97 | ENSGALG00000011485 | HCRT | Energy balance and potentially food intake | (17) |
| 1 | 6.03 | ENSGALG00000013422 | RHNO1 | Down-regulated in efficient residual feed intake birds | (18) |
| 7 | 5.8 | ENSGALG00000006217 | S100B | Involved in chicken skeletal muscle development | (19) |
| 11 | 5.84 | ENSGALG00000003273 | HSDL1 | Maintenance of secondary sexual characteristics and sex differentiation by metabolising hormones | (20) |
| 33 | 5.60 | ENSGALG00000047132 | LOC112529929 | keratin, type II cytoskeletal 4-like | - |
| 33 | 4.86 | ENSGALG00000042837 | LOC100858942 | glucagon-like | - |
| 11 | 4.68 | ENSGALG00000003201 | FBXL8 | Tumour suppressor in humans | (21) |
| 1 | 3.98 | ENSGALG00000011798 | HEBP1 | Heme-binding protein 1 | - |
| 1 | 3.98 | ENSGALG00000011799 | HEBP1 | Heme-binding protein 1 | - |
| 1 | 3.67 | ENSGALG00000025906 | HS6ST3 | heparan sulfate 6-O-sulfotransferase 3 | - |
| 33 | 3.60 | ENSGALG00000030629 | KRT6A | Associated with frizzle feather | (22) |
| 33 | 3.60 | ENSGALG00000035972 | KRT75 | Role in hair and nail formation and responsible for frizzle chicken condition | (23) |
| 33 | 3.26 | ENSGALG00000044875 | KRT75L4 | Frizzle chicken condition in Chinese indigenous chicken | (24) |
| 8 | 3.23 | ENSGALG00000027608 | PIGC | phosphatidylinositol glycan anchor biosynthesis class C | - |
| 1 | 3.19 | ENSGALG00000032645 | HIST1H2A4L3 | Major component of nucleosome. | - |
| 1 | 3.19 | ENSGALG00000052649 | HIST1H2B5 | Major component of nucleosome. | - |
